# Pufferfish (*Tetraodon cutcutia*) sampled from freshwater river serves an intermediate reservoir of sucrose nonfermenting variant of *Vibrio cholerae* PS-4

**DOI:** 10.1101/2021.10.27.466062

**Authors:** Lipika Das, Sushanta Deb, Eiji Arakawa, Shinji Yamasaki, Subrata K Das

## Abstract

This study describes the genomic characteristics of *Vibrio cholerae* strain PS-4, which fails to ferment sucrose on thiosulfate□citrate□bile salt□sucrose (TCBS) agar medium. This bacterium was isolated from skin mucus of a freshwater pufferfish. In order to understand the sucrose nonfermenting phenotype, the genome of the strain PS-4 was sequenced. The gene encoding the sucrose specific phosphotransferase system IIB (*suc*R) was absent, resulting for the defective sucrose fermenting phenotype. In contrast, genes encoding glucose-specific transport system IIB (*pts*G) and fructose specific transport system IIB (*fru*A) were present and showing acid production while growing with respective sugars. The overall genome relatedness indices (OGRI) such as *in silico* DDH, average nucleotide identity (ANI) and average amino acid identity (AAI) were above the threshold value, i.e., 70% and 95–96%, respectively. Phylogenomic analysis based on genome wide core genes and the non-recombinant core-genes, strain PS-4 clustered with *Vibrio cholerae* ATCC 14035^T^. Further, genes encoding *ctx*, *zot*, *ace*, *tcp* and *rfb* were absent. It showed hemolytic activity and reacted strongly to the R antibody. This is the first report on the existence of *Vibrio cholerae* from the pufferfish adds to our knowledge of the new ecological niche of this bacterium.

**IMPORTANCE:** *Vibrio cholerae*, the causative agent of cholera, is a natural inhabitant of aquatic environments. It ferments sucrose, producing characteristic yellow colonies on TCBS agar. *Vibrio cholerae* strain PS-4 described in this study was a sucrose nonfermenting variant associated with pufferfish skin and did not produce yellow colonies on TCBS agar. Genes encoding sucrose specific phosphotransferase system IIB (*suc*R) was absent. The observed phenotype in the characteristic metabolic pathway indicates niche specific adaptive evolution for this bacterium. Our study suggested that nonfermenting phenotype on TCBS agar may not be considered always for *Vibrio cholerae* species delineation.

## INTRODUCTION

Vibrios are ubiquitous and abundant in the aquatic environment including estuaries, marine coastal waters and sediments, and aquaculture practices worldwide (1, 2). Several studies have demonstrated that Vibrios are associated with aquatic animals (3, 4). Rapid growth with a short generation time, salt tolerance ability and biofilm forming capacities have acquired the adaptive abilities and physiological flexibilities in the genus Vibrio to survive and flourish in the diverse oligotrophic environment (5). *Vibrio* spp. are Gram-negative bacteria belonging to the class Gammaproteobacteria. This group of bacteria is chemoorganotrophic, mesophilic and usually motile rods. These are either pathogenic (6) or non-pathogenic (7). *V. cholerae* O1, O139, and non-O1/non-O139 are autochthonous to the aquatic environment. Nearly, 200 recognized serogroups of *V. cholerae* are known till date. Among them only two, O1 and O139, are associated with epidemics and global pandemics of cholera. However, many *V. cholerae* isolated from the aquatic environment acquired virulence genes or their homologs with low or no pathogenicity (8). Thus, emergence of pathogenic *Vibrio* strains in the environment as a result of a potential gene cradle where bacteria can exchange genetic elements (9). Serotypes O1 and O139 possess two main virulence genes, cholera toxin (CTX) and toxin-coregulated pilus (TCP), play a pivotal role in the pathogenicity of *V. cholerae* (10). However, other strains (non-O1/non-O139) that cause sporadic cases of diarrhea have toxigenic potential attributed to secretion systems (T3SS and T6SS) and other accessory toxin like zonula occludens toxin (Zot) (11). Additionally, they have other gene coding to hemolysin, helping to the colonization of the intestine (12). The absence of cholera enterotoxin was also reported in *V. cholerae* non-O1/non-O139 isolated from the water of several reservoirs. Some studies have demonstrated the antigenic translation of *V. cholerae* non-O1/non-O139 to *V. cholerae* O1 in favourable conditions (13). Whole genome sequence analysis is a widely accepted procedure for comprehensive evaluation of phylogenetic relations by identifying single nucleotide polymorphisms (SNPs) of *V. cholerae* pandemics. Besides, genomic data can be used for the characterization of endemic strains and estimation of *V. cholerae* transmission routes.

*Vibrio cholerae*, the causative agent of cholera, is a natural inhabitant of aquatic environments. *Vibrio* spp. are frequently isolated from fish, fish products, and edible shellfish, and a large number of species are pathogenic to different hosts. Recent evidence supports the hypothesis that fish may also be intermediate reservoirs and vectors of *V. cholerae* (14, 15). Indeed, cholera was associated with the consumption of fish and fish diet in different parts of the world (16, 17). In this study, we employed the culturable approach for the isolation of bacteria associated with skin mucus of fresh water pufferfish collected from Mahanadi River, Cuttack, India. Pufferfish belong to the order Tetraodontiformes and are omnivorous and have benthic habitat. They produces toxin causes physiological disorder in human (18). Report on the discovery of *V. cholerae* in the skin mucus of fresh water puffer fish has not been reported so far. Here, we described the biochemical characteristics, genomic analysis and virulence properties of *Vibrio cholerae* strain PS-4 isolated from fresh water pufferfish.

## RESULTS AND DISCUSSION

### Isolation and identification of pufferfish skin associated bacteria

Mucosal surfaces of fish is one of the nutrient-rich surfaces available to the aquatic microorganisms. In most cases, the bacteria were isolated straight way from the mucus layer without any enrichment, indicating the abundance of microbial communities of fish species (19). In this study, 26 bacterial strains were isolated from the Pufferfish skin. 16S rRNA gene sequence analysis identified these bacteria under twelve taxa belonged to the class Gammaproteobacteria, Betaproteobacteria, Bacilli and Flavobacteriia. Among them, majority of the strains were assigned to the class Gammaproteobacteria. Bacteria identified from the mucus layer of pufferfish represent the genus *Acinetobacter*, *Shewanella*, *Bacillus*, *Aeromonas*, *Serratia*, *Moraxella*, *Delftia*, *Staphylococcus*, *Chryseobacterium*, *Exiguobacterium*, *Chromobacterium and Vibrio*. All the strains were closely related to the respective bacterial taxa with a 16S rRNA sequence similarity of more than 98% **(Table S1).** It is known that mucosal surface of the skin and its associated microbiota provides protection against possible pathogens contributing to host immune maturity (20) and serves as a natural niche for aquatic mucosal pathogen evolution (15). Diversity and phylogenetic studies on Vibrio from various environmental and clinical samples have been investigated. However, presence of *Vibrio cholerae* species from the skin mucosal surfaces of pufferfish has not been reported so far (21). Like many other fish, no analysis of composition in bacterial species as well as evaluation of potential pathogenicity of the microbes associated in the skin mucosal surfaces of pufferfish and their distinction between potentially virulent vs non-virulent was performed. Thus, *Vibrio cholerae* strain PS-4 isolated from pufferfish was used for detail studies.

### Phenotype and serogroup of *Vibrio cholerae* strain PS-4

The cells of strain PS-4 were Gram negative, positive for oxidase and catalase. It showed a hemolytic activity on blood agar. Typically, *V. cholerae* ferments sucrose, producing characteristic yellow colonies on TCBS agar. In contrast, strain PS-4 was sucrose fermentation negative and produced green colonies on this medium. In addition, it showed yellow colonies on Luria Bertani Agar medium supplemented with either glucose or fructose similar to the *Vibrio cholerae* strain N16961 **(Fig.1).** Genome analysis of the strain PS-4 revealed that the gene encoding the phosphotransferase system (PTS) sucrose-specific IIB (*suc*R) component was absent, accounting for the defective sucrose-fermenting phenotype. In contrast, genes encoding glucose-specific transport system IIB (*pts*G) and fructose specific transport system IIB *fru*A) were present and showing acid production while growing in presence of respective sugars. Our study based on biochemical characterization and genomic analysis suggested that nonfermenting phenotype of *Vibrio cholerae* on TCBS agar may not be considered always for its species identification.

**Fig. 1.**
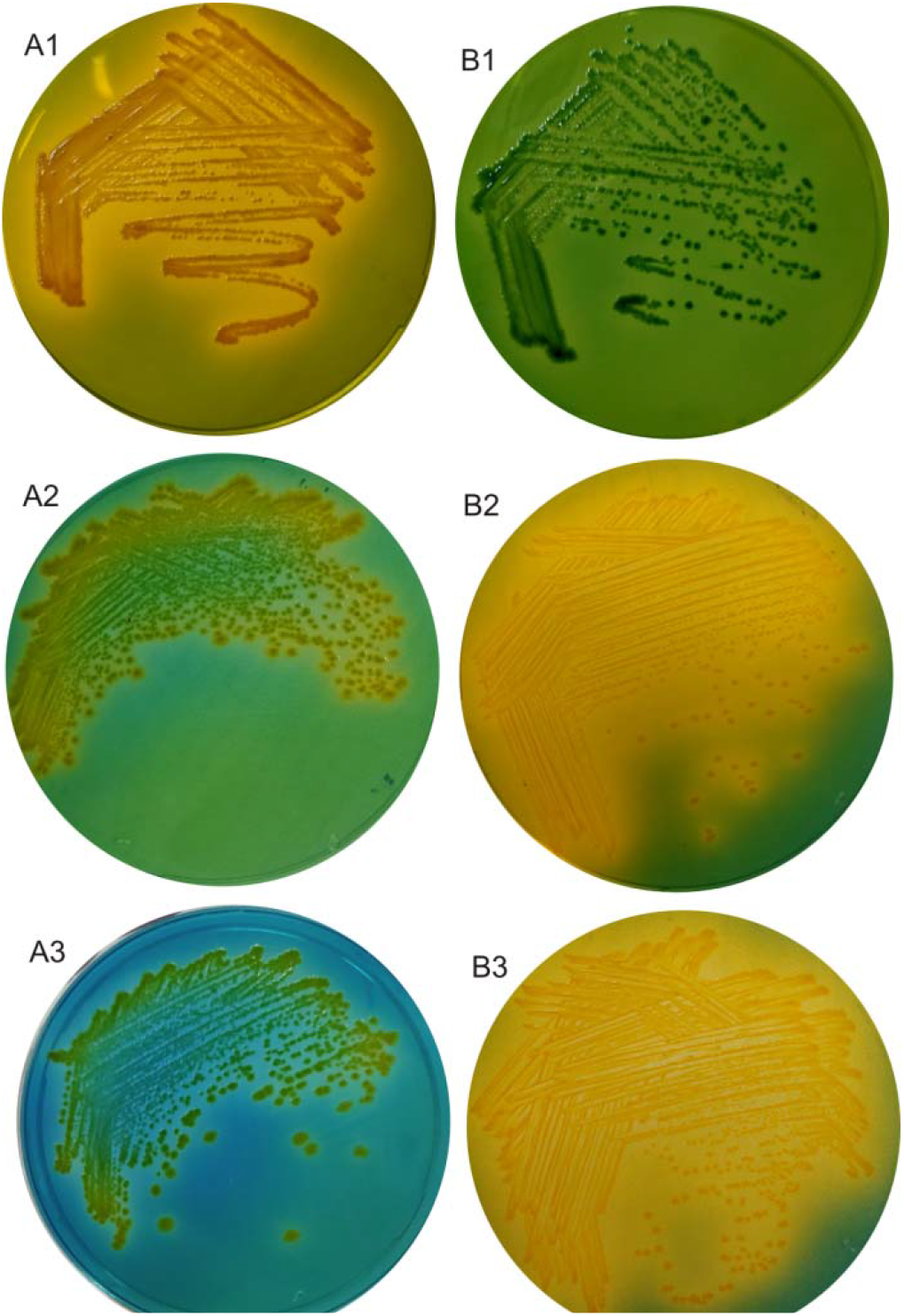
Growth response of *V. cholerae* strain N16961 and *V. cholerae* strain PS-4 on TCBS (Thiosulfate-citrate-bile salts-sucrose agar) and Luria Bertani agar supplemented with 0.5% glucose or 0.5% fructose and 2.0 mg/L bromothymol blue. *V. cholerae* N16961: A1, growth on TCBS; A2, growth on Luria Bertani agar plus glucose; A3, growth on Luria Bertani agar plus fructose. *V. cholerae* PS-4: B1, growth on TCBS; B2, growth on Luria Bertani agar plus glucose; B3, growth on Luria Bertani agar plus fructose.

The serotyping result showed that strain PS-4 reacted strongly to the R (rough) antibody. Each antiserum has absorbed with R antigen. Moreover, BLAST analysis of strain PS-4 scaffold sequences with the O antigen region of all O serogroups available in the NCBI database showed high homology with the part of the sequence of O127 antigen. Thus, it can be assumed that the phenotype of O antigen of strain PS-4 is R, but the genotype seems to be O127 **(Supplementary table 1).**

### Genomic features of *Vibrio cholerae* strain PS-4

The sequence of the *V. cholerae* strain PS-4 comprised two circular chromosomes, in which chromosome I contained 27,84,636 bp, while chromosome II contained 98,4931 bp. The overall GC content was of 47.61%. There are 3,364 predicted CDS, of which 3,304 had a homologous function predicted, and 205 were annotated as hypothetical proteins. The strain’s genome contained 104 tRNA genes and 31 rRNA genes. The predicted ORFs were further classified into COGs functional groups **(Fig.2)**

**Fig. 2.**
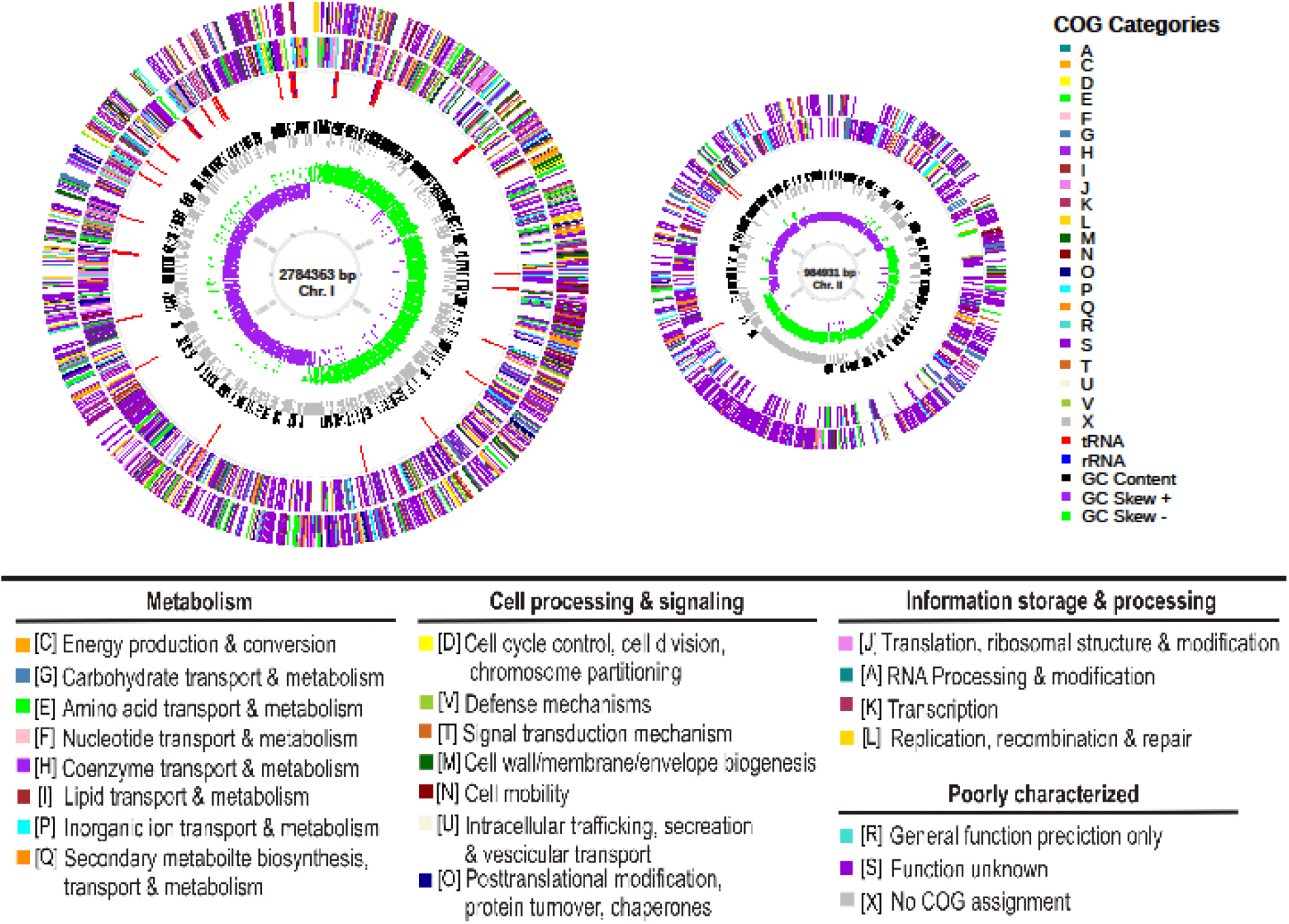
Circular graph of *Vibrio cholerae* strain PS-4 genome. Concentric outer to inner rings represents the protein coding genes on the forward strand, protein-coding genes on the reverse strand, tRNA (red) and rRNA (blue) genes, GC content, GC skew, scale marks of the genome. Protein-coding genes are color-coded according to their COG categories.

### Genome based analysis and phylogeny of *Vibrio cholerae* strain PS-4

Prokaryotic taxonomy provides a structure for proper identification of microorganisms. In this regard, we evaluated the DNA–DNA similarity, the average nucleotide identity (ANI), the average amino acid identity AAI values in combination with SNP-based phylogeny and genome wide core genes based phylogenetic analysis with the validly named type species to justify strain PS-4 belonging to *Vibrio cholerae*. The ANI and AAI values between strain PS-4 and the type species of *Vibrio cholerae* ATCC 14035 were higher than the threshold values (95-96%), justifying both strains belong to the same species (22). Further, *in silico* genomic DNA–DNA similarity value was higher than the 70 % cut off to define bacterial species (23). Thus, ANI, AAI and *in silico* DNA-DNA hybridization (*is*DDH) data indicated that the strain PS-4 belong to the same species of *Vibrio cholerae* **(Table 1).** SNP-based phylogeny revealed that strain PS-4 clustered with non-O1/non-0139 *Vibrio cholerae* strains **(Fig. 3)**. The maximum-likelihood (ML) phylogenetic tree based on genome wide core genes showed that strain PS-4 which clustered with *Vibrio cholerae* ATCC 14035 **(Fig. 4)** should be considered now as belonging to *Vibrio cholerae*. In addition, the non-recombinant core-genome based phylogenetic tree, strain PS-4 clustered with *Vibrio cholerae* ATCC 14035 in the same clade (**Fig. S1)** compared to the tree generated using core genomes to determine the taxonomic position (**Fig. 4**), indicating robustness of tree topology.

**Table 1.**
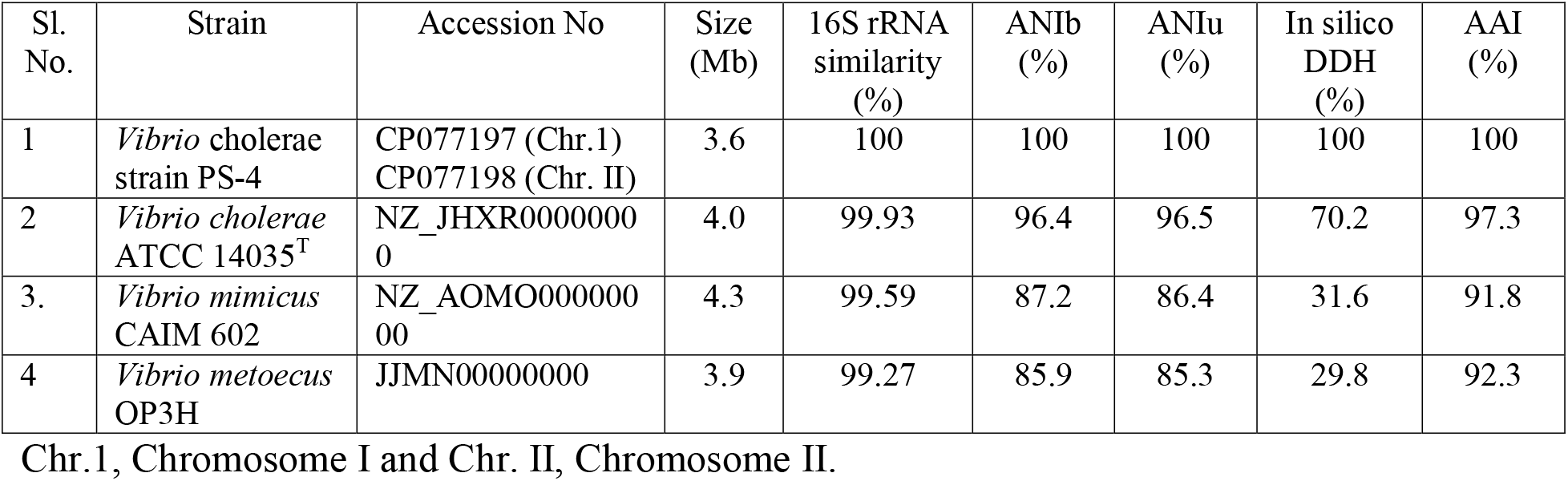
Comparison of the genomic characteristics with closely related species of Vibrio.

**Fig. 3.**
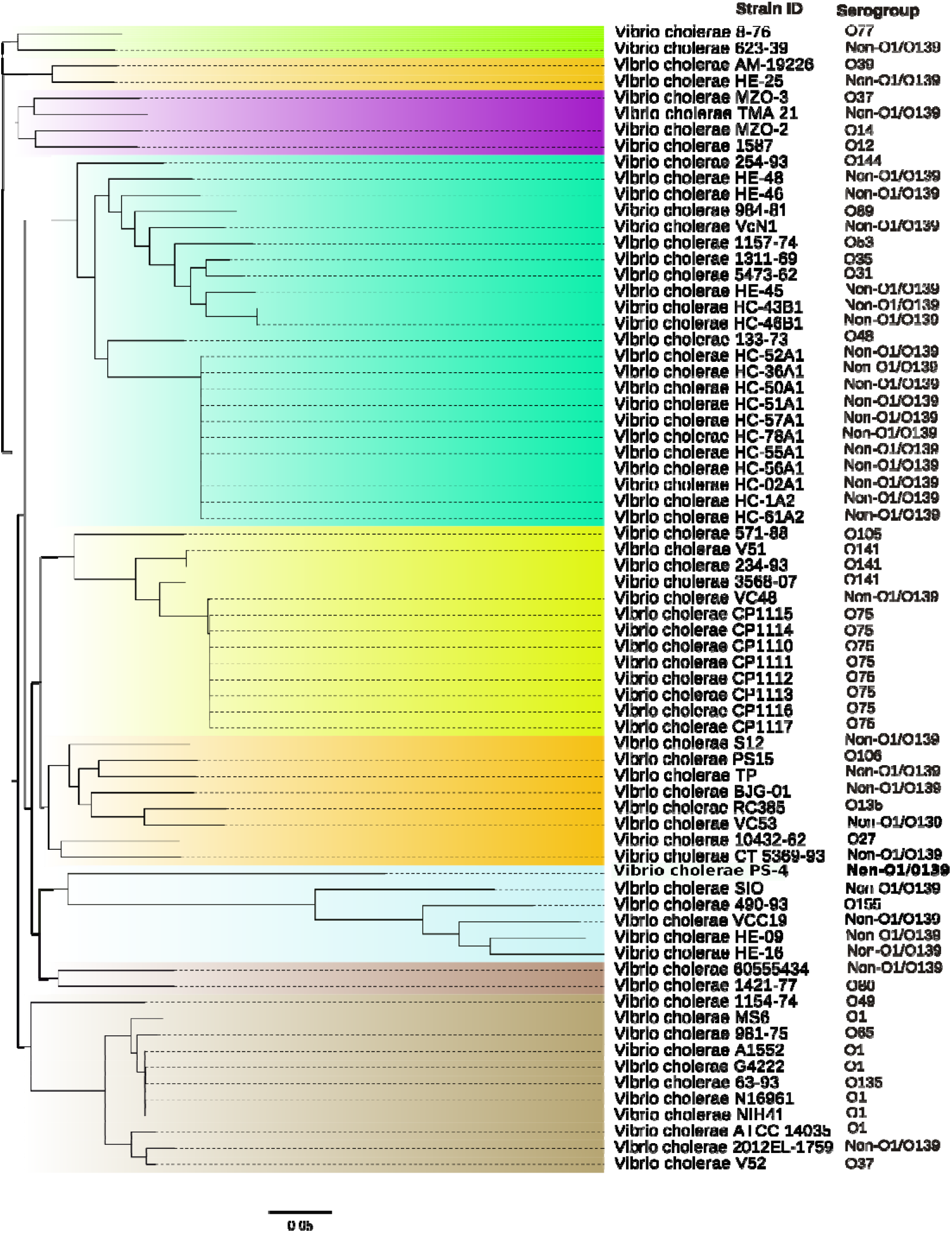
Maximum-likelihood phylogenetic tree based on genome-wide SNPs.

**Fig. 4.**
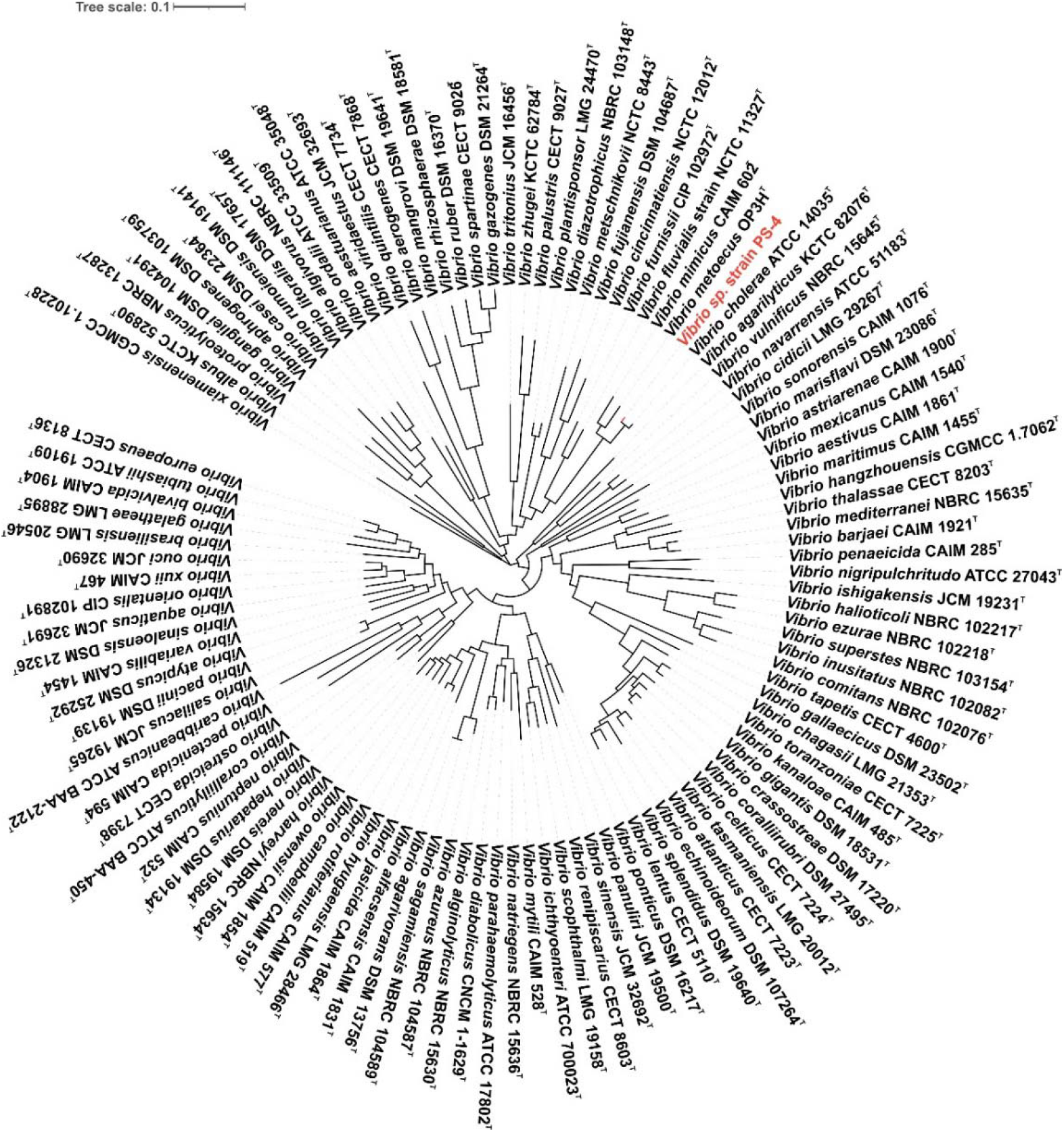
Core genome based phylogenetic tree based on the alignment of core genes from 131 type strains of Vibrio of all species with correct validly published names. Phylogenetic position of strain PS-4 high light in red color.

### Virulence properties

Virulence genes identified in the genome of strain PS-4 are listed in **Table-2**. Altogether 28 virulence associated genes were compared with 71 *Vibrio cholerae* strains. The gene sequences of individual gene were compared with the reference toxigenic *V. cholerae* O1 El Tor strain N16961. Hierarchical clustering analysis illustrated that strain PS-4 shared maximum sequence similarity with non-toxigenic *Vibrio* isolates viz., HE-16, HE-09, VCC19, SIO and 490 93 in a monophyletic clade (**Fig. 5**). It is known that *hly*A, responsible for hemolytic activity occasionally reported from non-toxigenic non O1/O139 serogroups (24, 25). The *hly*A gene of strain PS-4 showed 97% sequence similarity that of *V. cholerae* O1 El Tor strain N16961. However, other non-toxigenic strains (HE-16, HE-09, VCC19, SIO and 490 93) of the same clade were showing sequence divergence (<98% nucleotide identity). Earlier report suggested that most of the non-O1/non-O139 strains did not have the *ctx* (cholerae enterotoxin), *tcpA* (toxin-coregulated pilus), *zot* (zonula occludens toxin), *ace* (accessory cholera enterotoxin) and *rfb* (lipopolysaccharide biosynthesis) genes (10). Genome analysis revealed that *ctx*, *zot*, *ace*, *tcp* and *rfb* were absent in PS-4, hence this organism could be regarded as a non O1/O139 serogroup. In *V. cholerae* type VI secretion system plays a critical role to deliver toxins into adjacent target cells and compete against other bacteria with toxins that disrupt lipid membranes, cell walls and actin cytoskeletons (26). Type VI secretion system consist of a large number of virulence associated secretion (*vas*) genes and *vgr*G effector protein (27). In this regard, type VI secretion system of strain PS-4 encoded 15 genes. These genes showed sequence similarity more than 94% viz., *vas*L (97.86%), *vip*A (97.83%), *vas*G (99.15%), *vas*D (99.37%), *vas*A (99.09%), *vas*I (97.71%), *vas*K (97.57%), *vas*F (97.02%), *vas*J (98.29%), *vas*C (98.85%), *vas*B (98.42%), *vas*H (98.43%), *vas*E (98.05%), *vgr*G2 (97.53%) and *vgr*G3 (94.49%) with *V. cholerae* O1 El Tor strain N16961. Besides, the gene encoding the thermolabile hemolysin (*tlh*) is also considered a signature molecular marker for the species (28). This gene is rarely reported from nonclinical strains. DNA sequence of *tlh* identified in the strain PS-4 showed 60% similarity with *Vibrio parahaemolyticus*. Thus, existence of *Vibrio cholerae* strain from the pufferfish skin adds to our knowledge of the new ecological niche of this bacterium.

**Table 2.**
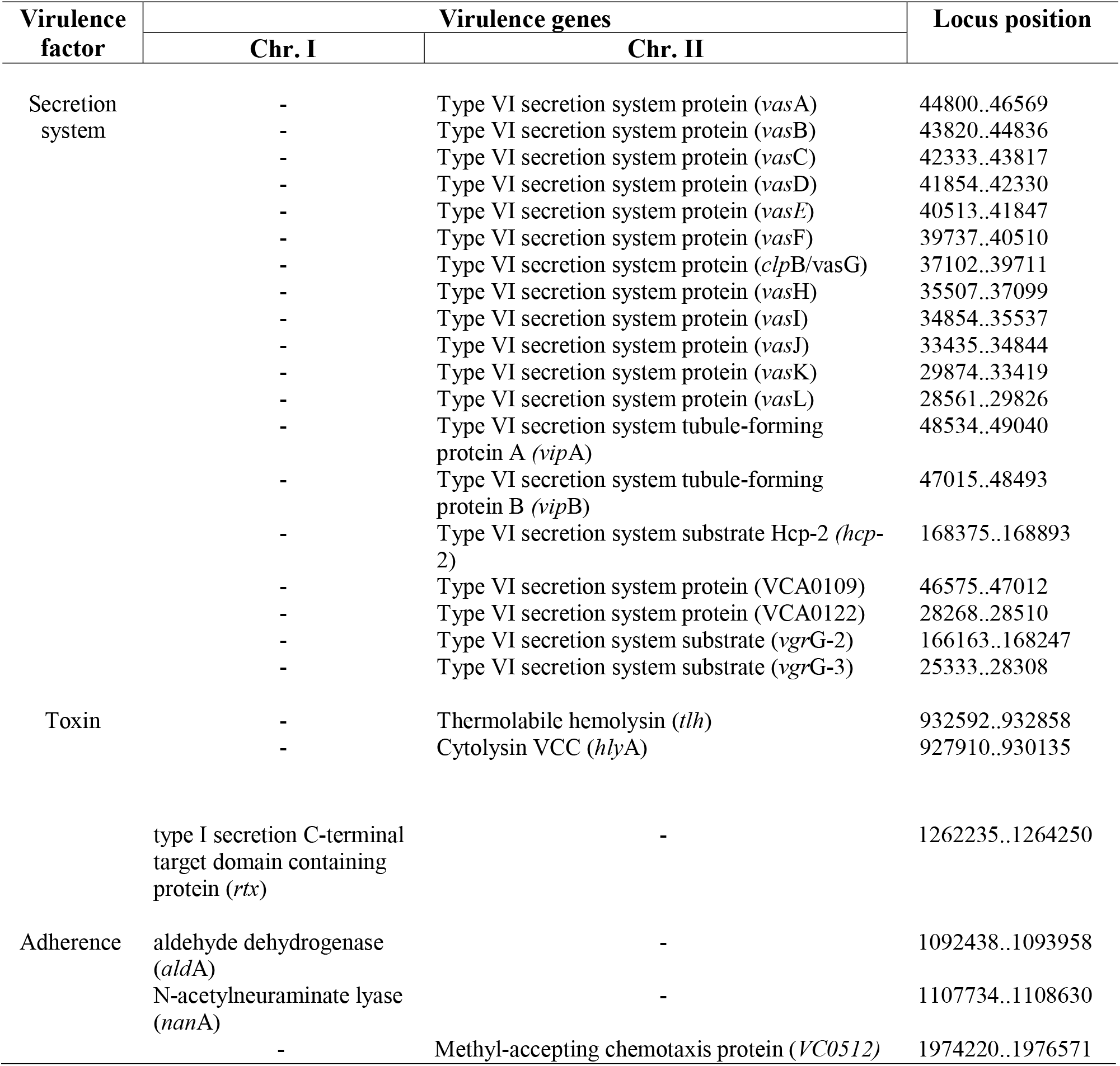
Virulence factors of *Vibrio cholerae* strain PS-4. -, not detected.

**Fig. 5.**
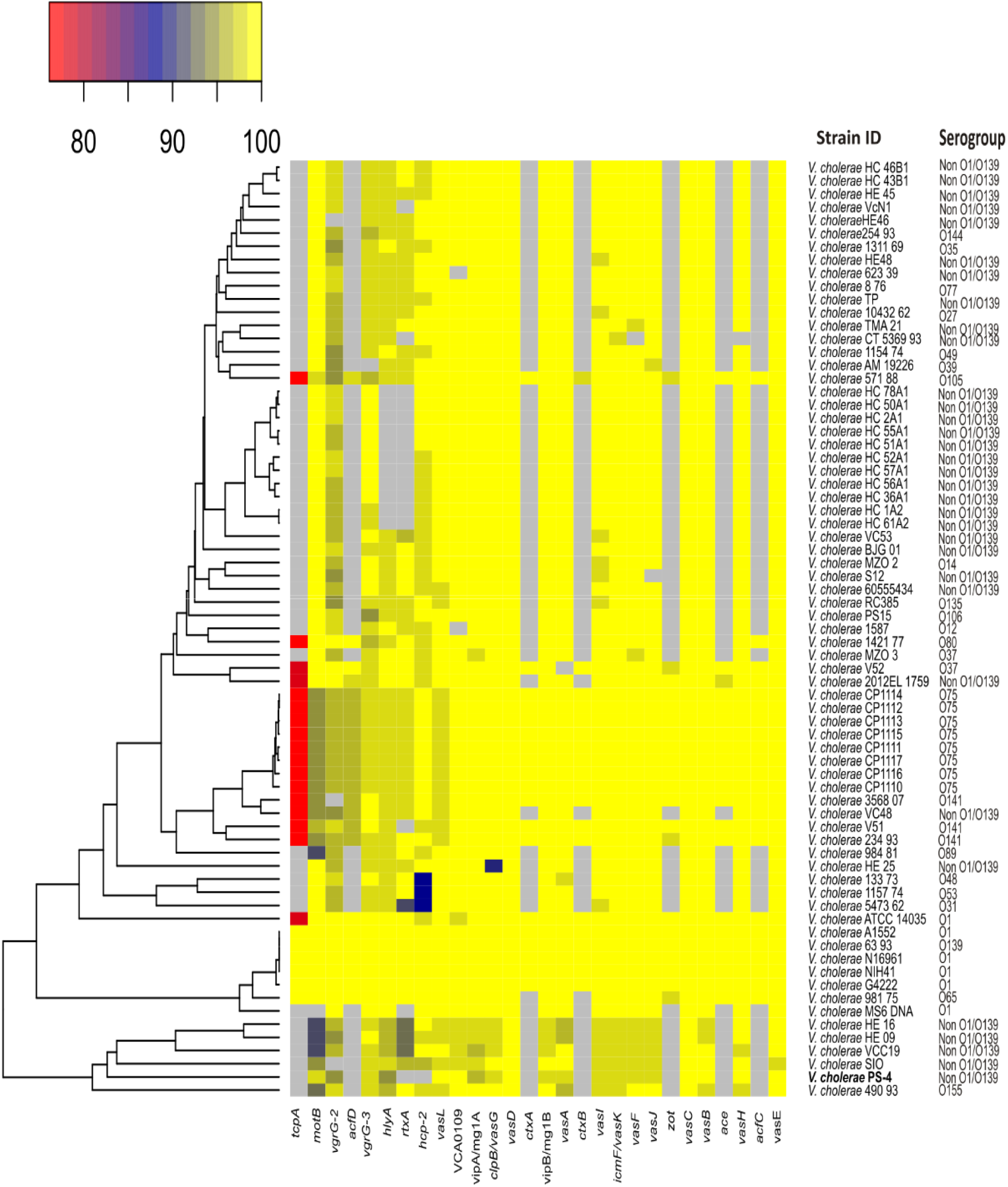
Conservation in nucleotide sequences of orthologous virulence genes in *Vibrio cholerae* strain PS-4 with reference strains. Top bar represents percent nucleotide sequence identity. Gray boxes shows missing genes. The hierarchical clustering of the strains was based on average linkage method and Manhattan distance similarity metric.

## MATERIALS AND METHODS

### Bacterial strain and growth medium

The pufferfish (*Tetraodon cutcutia*) samples were collected from Mahanadi River, India (coordinates: 20°26’46.6”N 85°44’28.3”E) in the month of August, 2018 and transported to the laboratory in a plastic container with river water. For isolation of bacteria, mucus on the skin of pufferfish were taken using sterile cotton swabs and transferred into 1 ml sterile phosphate buffer saline (PBS), pH 7.4. The bacteria from the cotton swabs were suspended in PBS by vigorous vortexing and the suspension was used as a master mix (29). An aliquot of (100 μl) master mix sample was serially diluted using PBS and plated onto nutrient agar (BD, Difco). All plates were incubated at 30 °C corresponding to the river water temperature for 2 days. Several colonies that developed at 30 ^o^C were picked and purified by repeated streaking on the same medium. Cultures were maintained on nutrient agar (BD, Difco) and stored at 4°C for short-term preservation. For long-term preservation the culture were stored at −80°C in 15% (v/v) glycerol.

### Phenotypic features and identification of serogroup of *V. cholerae* strain PS-4

Gram staining was carried out using the commercial kit (Becton-Dickinson, USA). Oxidase activity was assayed with discs impregnated with dimethyl p-phenylenediamine (Hi-Media, India). Catalase activity was assayed by mixing a pellet of a freshly centrifuged culture with a drop of hydrogen peroxide (10 %, v/v). Growth and reaction to ferment sucrose was tested on thiosulfate□citrate□bile salt□sucrose (TCBS) agar medium (BD, Difco). Utilization of sugars were tested separately by adding 0.5% concentration of glucose or fructose in Luria Bertani Agar medium (BD, Difco) containing bromothymol blue 2.0 mg /L as pH indicator, at 37 °C for 48 h. To ascertain the haemolytic activity, strain PS-4 was streaked on Columbia blood agar base supplemented with 5% (v/v) defibrinated sheep blood followed by incubation at 37 °C for 48 h (29). Preparation of O-antisera and slide agglutination were performed as previously described (30).

### Identification of bacteria by 16S rRNA sequencing

Genomic DNA was extracted following the methods of Sambrook and Russel (31) and PCR was carried out using the bacterial universal primers 27F (5’-GAGTTTGATCCTGGCTCAG-3’) and 1525R (5’-AAAGGAGGTGATCCAGCC-3’) (32). The PCR product was purified using QIAQuick gel extraction kit (Qiagen) and sequenced in a capillary DNA analyzer (model number 3500, Applied Biosystems) following the manufacturer protocol. The 16S rRNA gene sequences were assembled using the sequence alignment editor program BioEdit (33) and compared with those in GenBank after BLAST searches (34) and using the EzBioCloud Database (35).

### Whole genome sequencing and annotation

The genomic DNA of *Vibrio cholerae* strain PS-4 was isolated using standard methods by Sambrook and Russel (31). DNA concentration and quality was measured using NanoDrop 8000 Spectrophotometer (Thermo Scientific). Combination of both short-read Illumina and long-read Oxford Nanopore sequencing platform was used to generate the high-quality complete genome sequence of *Vibrio cholerae* strain PS-4. Illumina short-read DNA sequencing was carried out as described earlier (29). For long read Nanopore sequencing, genomic library was prepared using the Nanopore Ligation Sequencing Kit (SQK-LSK109; Oxford Nanopore, Oxford, UK). The library was sequenced with an R9.4.1 MinION flow cell (FlO-MIN106) using MinKNOW v2.0 with the default settings. Barcode and adapter sequences from Nanopore long reads were trimmed using Porechop v0.2. (https://github.com/rrwick/Porechop) and reads with minimum of 1 kb length were filtered using seqtk v1.2 (https://github.com/lh3/seqtk) for downstream analysis. The hybrid genome assembly was performed using Unicycler version 0.4.9 (36) in hybrid assembly mode. The highly accurate illumina short reads were aligned against the long Nanopore reads to sort out random sequencing errors (36). The assembled genomes were annotated using the NCBI Prokaryotic Genome Annotation Pipeline (PGAP version 4.9) with default parameters (37). Completeness and contamination of the whole genome sequence were measured using CheckM (38). Genomic G+C content and assembly statistics were determined using Perl script (https://github.com/tomdeman-bio/Sequence-scripts/blob/master/calc_N50_GC_genomesize.pl).

### Comparative genomics

Several bioinformatics analysis tools were used to compare the genomic relatedness of strain PS-4 with reference genomes of validly published 131 type strains of *Vibrio* available in the National Centre for Biotechnology Information (NCBI) database (last accessed March 25, 2021). With the advent of next-generation sequencing and bioinformatics tools made possible to compare genomic data by *in silico* DDH (*is*DDH), average nucleotide identity (ANI) and average amino acid identity (AAI) values. The average nucleotide identity (ANI) was calculated using the python module pyani (https://github.com/widdowquinn/pyani) with ANIb method. In silico DDH similarity was measured with the help of Genome-to-Genome Distance calculator (formula 3) (23). Average amino acid identity (AAI) was estimated using the ‘aai_wf’ function implemented in compareM program (https://github.com/dparks1134/CompareM).

### Genome-wide SNPs determination and phylogenetic analysis

For SNP based phylogenetic analysis 70 complete or draft genome sequences of *V. cholerae* strains were retrieved from the NCBI database. Single-nucleotide polymorphisms (SNPs) were identified from genome assemblies using *V. cholerae* strain N16961 as reference for alignment using Snippy version v4.6.0 (https://github.com/tseemann/snippy). The recombinant region were removed using Gubbins version 2.3.4 (39) with default parameters. Core SNPs were extracted with the help of SNP sites (40) and a maximum-likelihood (ML) phylogenetic tree was constructed using RAxML version 8.2.4 (41) with GTRGAMMA model (42) for nucleotide substitution with gamma-distributed rate heterogeneity.

In addition, the use of whole genome sequences has been regarded as a promising avenue to determine the phylogenetic position of microorganisms. Analysis of evolutionary phylogeny based on core-genomes is the current gold standard for strain identification, which is superior to those based on a single gene marker or concatenated sequences of a few genes. Therefore, we performed the phylogenomic analysis based on genome wide core genes of available whole genome sequence of 131 type strains of all species with correct validly published names of Vibrio with more than 95% genome completeness. The genome sequence of the type strains were retrieved from the NCBI database (https://github.com/kblin/ncbi-genome-download/). The core genes extracted by UBCG pipeline (43) was concatenated and a maximum-likelihood tree reconstructed with the GTR model using RAxML tool (44). Further, the non-recombinant core-genome based phylogenetic tree was constructed following the method of Mateo-Estrada et al. (45).

### Comparative analysis of virulence genes

Virulence-associated proteins of strain PS-4 were identified using blastp program against the VFDB database (46) following the parameters (identity cutoff 75%, coverage cutoff 70%, E-value cutoff 1e-5). The virulence related genes of strain PS-4 were compared with the O1/O139 type of *Vibrio cholerae* and non-O1/non-O139 *V. cholerae* serogroup strains using the blastn algorithm (47). Heatmap was generated from nucleotide percentage identity by means of manhattan distance and average clustering method using heatmap2 function of gplots package (48) in R program (49).

### Nucleotide sequence accession number

The GenBank/EMBL/DDBJ accession number for the genome and 16S rRNA gene sequences of *Vibrio cholerae* strain PS-4 are CP077197 (Chromosome-1), CP077198 (Chromosome-2) and MW926953, respectively.

## Supporting information

DNA sequence alignment data

## ACKNOWLEDGMENTS

The author LD and SD acknowledges the Council of Scientific and Industrial Research (CSIR), New Delhi, Government of India and Institute of Life Sciences, Bhubaneswar for providing the research fellowship. This work was supported by the funding received from the Department of Biotechnology, Government of India (D.O.No. BT/BI/04/058/2002 VOL-II) to SKD under the Distributed Information Sub-Center (DISC) project.

## ETHICAL APPROVAL

This study was carried out with approval from the Institutional Animal Ethics Committee (Letter No. V-311-MISC/2017-18/ILS/884).

## AUTHOR CONTRIBUTIONS

S.K.D. developed the concept and design the experiments. S.K.D, E.A and S.Y. coordinated the experiments and analyzed the data. L.D., S.D. participated in laboratory experiments. L.D, S.D and S.K.D. wrote the manuscript. All of us read and approved the final manuscript. The authors declare no conflicts of interest.

