## Supplementary data, table 1 and figure S1 for "Pufferfish (*Tetraodon cutcutia*) sampled from freshwater river serves an intermediate reservoir of sucrose nonfermenting variant of *Vibrio cholerae* PS-4"

**Running title:** *Vibrio cholerae* from fresh water pufferfish skin


**Authors:** Lipika Das[1, 2], Sushanta Deb[1], Eiji Arakawa[3], Shinji Yamasaki[4] and Subrata K Das[1, 2*]

**Addresses:** [1]Institute of Life Sciences, Department of Biotechnology, Nalco Square, Bhubaneswar - 751023, India; [2]Regional Center of Biotechnology, NCR Biotech Science Cluster, 3rd Milestone, Faridabad, Haryana (NCR Delhi), India; [3]Department of Bacteriology I, National Institute of Infectious Diseases, Tokyo 162-8640, JAPAN and [4]Department of Veterinary Science, Graduate School of Life and Environmental Sciences, Osaka Prefecture University, Osaka 598-8531, Japan

*Corresponding author. Mailing address:
Institute of Life Sciences
Department of Biotechnology
Nalco Square, Bhubaneswar 751023 India

Phone: (+91) 674 230 4328
Fax: (+91) 674 230 0728

### Supplementary table

**Table S1:** Bacterial strain isolated from puffer fish skin

| Strain | Sequence length (bp) | GenBank accession number | Most closely related hit in EzTaxon server | Similarity (%) |
|---|---|---|---|---|
| PS-1 | 1532 | MW114828 | *Acinetobacter kanungonis* JCM 34131$^{T}$ | 100 |
| PS-2 | 1413 | MK165131 | *Shewanella xiamenensis* S4$^{T}$ (FJ589031) | 98.73 |
| PS-3 | 1425 | MK165132 | *Bacillus marisflavi* JCM 11544$^{T}$ (AF483624) | 99.86 |
| PS-4 | 1539 | MW926953 | *Vibrio cholerae* ATCC 14035 (X76337) | 99.93 |
| PS-5 | 1443 | MK165134 | *Exiguobacterium indicum* HHS31$^{T}$ (AJ846291) | 99.86 |
| PS-6 | 1425 | MK165135 | *Aeromonas veronii* CECT 4257$^{T}$ (X60414) | 100 |
| PS-7 | 1434 | MK165136 | *Serratia rubidaea* JCM 1240$^{T}$ (AB004751) | 99.79 |
| PS-8 | 1366 | MK165137 | *Chryseobacterium gambrini* DSM 18014$^{T}$ (AM232810) | 99.12 |
| PS-9 | 1429 | MK165138 | *Exiguobacterium indicum* HHS31$^{T}$ (AJ846291) | 99.93 |
| PS-10 | 1416 | MK165139 | *Bacillus aryabhattai* B8W22$^{T}$ (EF114313) | 100 |
| PS-11 | 1386 | MK165140 | *Delftia litopenaei* wsw-7$^{T}$ (GU721027) | 99.71 |
| PS-12 | 1370 | MK165141 | *Bacillus indicus* LMG 22858$^{T}$ (AJ583158) | 99.93 |
| PS-13 | 1413 | MK165142 | *Acinetobacter baumannii* ATCC 19606$^{T}$ (X81660) | 99.93 |
| PS-14 | 1421 | MK165143 | *Staphylococcus haemolyticus* MTCC 3383$^{T}$ (X66100) | 100 |
| PS-15 | 1423 | MK165144 | *Acinetobacter junii* CIP 64.5$^{T}$ (Z93438) | 99.58 |
| PS-16 | 1430 | MK165145 | *Aeromonas enteropelogenes* CECT 4487$^{T}$ (EU306837) | 99.86 |
| PS-17 | 1406 | MK165146 | *Shewanella xiamenensis* S4$^{T}$ (FJ589031) | 98.65 |
| PS-18 | 1439 | MK165147 | *Bacillus cereus* ATCC 14579$^{T}$ (AE016877) | 100 |
| PS-19 | 1424 | MK165148 | *Shewanella xiamenensis* S4$^{T}$ (FJ589031) | 98.74 |
| PS-20 | 1425 | MK165149 | *Aeromonas veronii* CECT 4257$^{T}$ (X60414) | 99.86 |
| PS-21 | 1419 | MK165150 | *Acinetobacter variabilis* NIPH 2171$^{T}$ (KB850112) | 99.58 |
| PS-22 | 1417 | MK165151 | *Moraxella osloensis* CCUG 350$^{T}$ (AB643599) | 99.08 |
| PS-23 | 1445 | MK165152 | *Exiguobacterium indicum* HHS13$^{T}$ (JF893462) | 99.93 |
| PS-24 | 1402 | MK165153 | *Chromobacterium alkanivorans* IITR-71$^{T}$ (JN210566) | 100 |
| PS-26 | 1422 | MK165154 | *Bacillus altitudinis* 41KF2b$^{T}$ (AJ831842) | 99.93 |
| PS-27 | 1448 | MK165155 | *Exiguobacterium indicum* HHS31$^{T}$ (AJ846291) | 99.86 |

**Supplementary figure**

Figure S1. Non-recombinant core-genome based phylogenetic tree from 131 type strains of Vibrio of all species with correct validly published names. The phylogenetic position of PS-4 is highlighted in colour

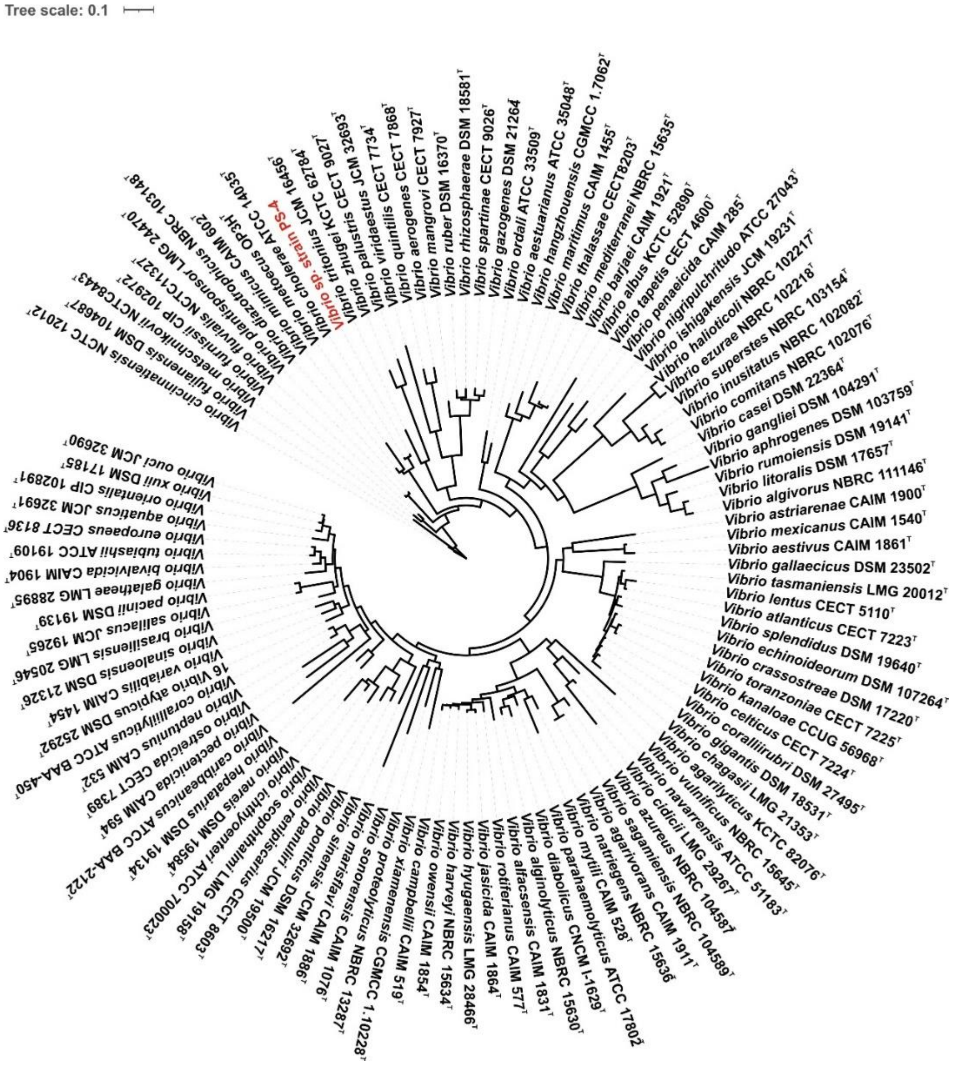


**Figure S1**
