## Supplementary figures and images for "Pufferfish (*Tetraodon cutcutia*) sampled from freshwater river serves an intermediate reservoir of sucrose nonfermenting variant of *Vibrio cholerae* PS-4"

### Supplementary table 2

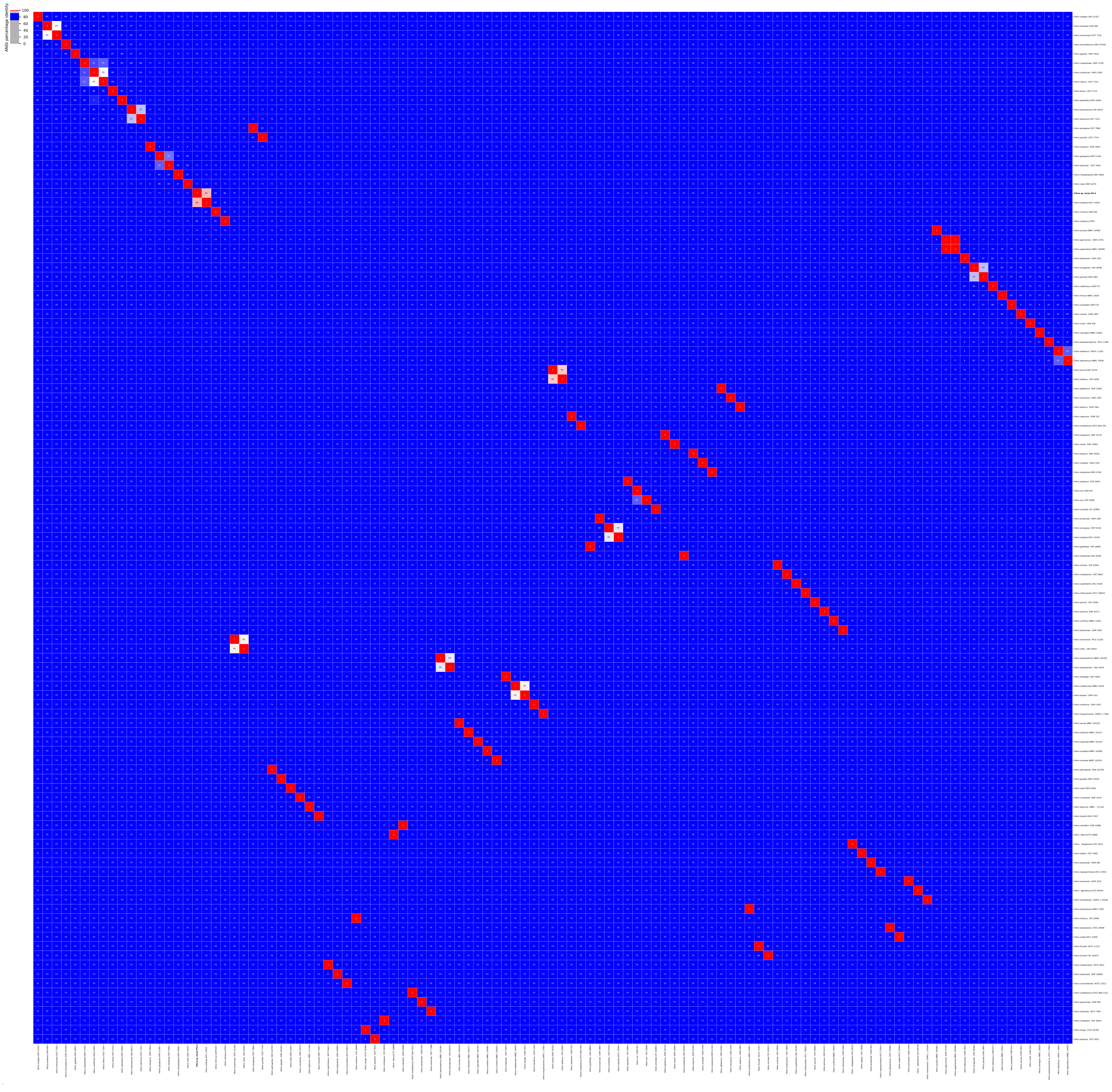
